# Changes in biomarkers of metabolic stress during late gestation of dairy cows associated with colostrum volume and immunoglobulin content

**DOI:** 10.1101/2022.07.18.500308

**Authors:** Renato M. Rossi, Faith M. Cullens, Paola Bacigalupo, Lorraine M. Sordillo, Angel Abuelo

## Abstract

The objective of this observational study was to compare the metabolic status of dairy cows during the last 6 wk of gestation based on colostrum volume and Ig content. For this, healthy Holstein cows were randomly selected from 3 commercial herds in Michigan. In each farm, four cohorts of 21 cows (one per season), stratified by parity, were enrolled (n=228). Cows were blood sampled weekly during the last 6 wk of gestation, and biomarkers related to nutrient utilization, oxidant status, and inflammation were quantified in serum. Cows were milked within 6 h of calving and the volume of colostrum produced was recorded and an aliquot collected. Concentration of IgG, IgA, and IgM were measured by radial immunodiffusion. Cows were grouped into high colostrum producer (**HCP**) or low (**LCP**), high IgG (**HIG**) or low (**LIG**), high IgA (**HIA**) or low (**LIA**), and high IgM (**HIM**) or low (**LIM**). For volume category, we arbitrarily defined 6 L of colostrum (4 L for first and 2 L for second feeding of calves) as the cut-off point, whereas for IgG we used the industry standard of ≥ 50g/L. To create groups of low and high IgM or IgA, we used the median of these Ig as the cutoff point. Colostrum volume was lowest in winter, but no differences were observed among parity groups. Conversely, colostrum IgG concentration was highest in fall and winter, but colostrum IgM was lowest at these seasons. However, colostrum Ig content only showed a negative weak correlation with volume (Spearman rho < -0.28). Compared to LCP, HCP cows had higher concentrations of antioxidant potential, BHB and lower cholesterol and oxidant status index. HIG cows showed higher concentrations of glucose compared to LIG. HIA cows had higher concentrations of cholesterol, reactive oxygen and nitrogen species, oxidant status index, and total protein, while BHB, and glucose were lower compared with LIA. Biomarkers of metabolic stress were not significantly different between HIM and LIM. Nevertheless, the differences observed did not result in differences in inflammatory status between animals in any of the colostrum variable categories analyzed, suggesting that physiological homeostasis was not disrupted during late gestation in association with the colostrum variables studied. Overall, the great variability observed in colostrum variables suggests that colostrogenesis is a complex and multifactorial process. However, our results suggest that greater availability of antioxidants during late gestation could support the production of higher volumes of colostrum, which needs to be explored in future trials.

**INTERPRETIVE SUMMARY:** **Changes in biomarkers of metabolic stress during late gestation of dairy cows associated with colostrum volume and immunoglobulins content.** *By Rossi et al*., *page XXXX*. We investigated associations between metabolic stress during the last 6 wk of gestation and the volume and immunoglobulin content of the colostrum produced. We observed that cows producing more than 6 L of colostrum exhibited increased metabolic activity during late gestation. Also, a greater blood antioxidant activity throughout late gestation was observed in cows with higher yields of colostrum, suggesting that greater availability of antioxidants might support the production of higher volumes of colostrum. Therefore, further studies should evaluate whether supplementation with additional antioxidants supplement during late gestation can improve colostrum yield.

## INTRODUCTION

Despite improvements in dairy calf health management practices over the last decades, preweaning morbidity and mortality incidence risks in US herds are still high, at 33.9% and 5%, respectively (Urie et al., 2018). One of the major contributors to preweaning disease occurrence is failure of transfer passive immunity, a problem that still has a high prevalence of 13% in US dairy herds (Raboisson et al., 2016). Providing insufficient volume and/or low quality of colostrum within the first hours of life is the major contributing factor to failed immunity transfer in calves (Morin et al., 1997). Traditionally, a volume of colostrum of 10 – 12% of the calves’ body weight, given in one single meal, was recommended as a feeding strategy to transfer passive immunity successfully (Godden, 2008). However, feeding 4 L of high-quality colostrum within 6 hours of birth and 2 L at 12 h of life has been recently recommended to optimize the transfer of passive immunity and calf health (Hammon et al., 2013; Godden et al., 2019). In fact, calves that received a second colostrum meal within the first 12 h of birth had greater ADG and lower failed immunity transfer and preweaning morbidity risks than calves that only received 1 meal (Abuelo et al., 2021).

However, there is considerable variability in colostrum production among cows (Morin et al., 1997; Gavin et al., 2018; Kessler et al., 2020), making it difficult to harvest the volume of colostrum needed to sustain this feeding regime in some cases. Colostrogenesis has been an active area of research and some aspects, such as IgG transfer from bloodstream, have been reviewed extensively (Barrington et al., 2001; Baumrucker and Bruckmaier, 2014). However, the scientific evidence on factors affecting the volume of colostrum being produced is still limited. Colostrogenesis starts 3 to 4 weeks before calving. At this time, cows start experiencing metabolic adaptations in preparation to the onset of lactation. However, cows may develop metabolic stress if they fail to physiologically adapt to the profound increase in nutrient requirements associated with fetal growth and milk production (Sordillo and Mavangira, 2014). Metabolic stress is characterized by excessive lipid mobilization, oxidative stress, and inflammatory dysfunction (Abuelo et al., 2015). The negative effect of metabolic stress on the immune function, health, and production of dairy cattle during this period is well established (Kehrli Jr et al., 1989; Sordillo and Aitken, 2009; Bradford et al., 2015). To the best of our knowledge, however, the association between metabolic stress biomarkers and colostrum production have not yet been examined. Colostrum is more concentrated in nutrients than milk (Godden, 2008), which might result in important nutritional demands for the cow. We, therefore, hypothesized that cows producing high volumes of colostrum and quality as assessed by immunoglobulin content would exhibit increased metabolic activity during late gestation. Thus, the objective of this observational study was to identify the association between biomarkers of metabolic stress during late gestation and colostrum volume and concentration of IgG, IgA, and IgM.

## MATERIALS AND METHODS

### Animals, Feed, Farms, and Management

All procedures were approved by the Michigan State University Institutional Animal Care and Use Committee (protocol 04/18-065-00) and animals were enrolled with owners’ consent. This prospective cohort study was conducted using a convenience sample of three commercial Michigan dairy farms, selected based on location within 50 miles to the university and willingness to participate in the study. The study was designed to have one cohort per season at each farm, a total of four cohorts, to account for the documented changes in colostrum yield associated with season (Gavin et al., 2018; Borchardt et al., 2022). Sampling occurred during the period between June 2019 and September 2020.

The sample size was calculated using an online calculator (https://epitools.ausvet.com.au) to achieve 90% confidence and 5% precision of within-herd prevalence, resulting in 21 animals per cohort per farm (n=252). Healthy Holstein cows were selected using randomization software (https://www.graphpad.com/quickcalcs/randomSelect1/) among those expected to calve 6-8 weeks after the start of sampling from a list of cows generated by the herd management software. Healthy animals were defined as not being in the sick pen, not currently receiving any medical treatment, and not displaying sick cow behavior based on the subjective interpretation of the research staff. To reflect common demographics of dairy farms, random selection was stratified by parity groups of cows entering their first, second, or third to fifth lactation. Exclusion criteria for enrollment were cows with a successful breeding later than 150 DIM, and body condition score (BCS) under 2 or over 4 on a scale of 1 – 5 (Wildman et al., 1982). Finally, data from 24 enrolled cows were excluded from analyses due to abortions, injuries resulting in euthanasia, deaths prior to calving, or failure to obtain colostrum data (samples not collected or yield not recorded). Therefore, the complete data from 228 cows were included in the analyses.

Housing and characteristics of management practices of late-gestation cows of the three farms are reported in Table 1. Cows had ad libitum access to a total mixed ration and water for the entire dry cow period. Farms A and B had two dietary groups for dry cows (far-off and close-up) whereas farm C managed all dry cows in the same diet. Farms A and C had separate groups for heifers and multiparous cows, whereas in farm B heifers and multiparous cows were separated in the far-off group but not the close-up. Diets were formulated by the farms’ nutritionist to meet or exceed NRC (2001) recommendations. Samples of all total mixed rations were collected at 2-weeks intervals from the feed bunk at the time of distribution and sent to an external laboratory for chemical composition analysis (Cumberland Valley Analytical Services, Waynesboro, PA). The composition and chemical analysis results of the diets are summarized in Supplementary Tables S1 and S2.

**Table 1:** Description of herd, housing, and management practices for late-gestation cows in the study farms.

|  | Farm A |  |  |  | Farm B |  |  | Farm C |  |  |
| --- | --- | --- | --- | --- | --- | --- | --- | --- | --- | --- |
|  | Close up cows | Close up heifers | Far-off cows | Far-off heifers | Close up cows / heifers | Far-off cows | Far-off heifers | Close-up cows / heifers | Far-off cows | Far-off heifers |
| Pen design | Free stall | Free stall | Free stall | Free stall | Bedded pack | Free stall | Free stall | Bedded pack | Free stall | Free stall |
| Heat abatement | 137 cm diameter fans on stall area |  | None | None | 91 cm diameter fans on bedding areas |  | None | 61 cm diameter fans on bedding areas |  |  |
| Target dry period length (d) | 60 |  |  |  | 60 |  |  | 45 |  |  |
| Target duration of close-up diet | 21 |  |  |  | 21 |  |  | Same diet for whole dry period |  |  |
| Rolling yearly milk yield per cow (kg) | 15,049 |  |  |  | 14,508 |  |  | 14,011 |  |  |
| Rolling yearly average of cows in milk (n) | 3,712 |  |  |  | 1,054 |  |  | 1,608 |  |  |
| Min-Max ambient temperature (°C)<sup>1</sup> |  |  |  |  |  |  |  |  |  |  |
| *Summer* | 17.2 to 30.1 |  |  |  | 17.8 to 29.4 |  |  | 15.2 to 30.5 |  |  |
| *Fall* | 5.5 to 14.8 |  |  |  | 5.5 to 15.1 |  |  | 11.1 to 22.2 |  |  |
| *Winter* | -6.8 to 1.0 |  |  |  | -6.7 to 1.1 |  |  | -6.1 to 2.8 |  |  |
| *Spring* | 1.3 to 11.0 |  |  |  | 1.1 to 12.2 |  |  | 0.8 to 12.8 |  |  |

<sup>1</sup> Data from the National Environmental Satellite, Data, and Information Service station closest to each farm ([https://www.ncei.noaa.gov/access/past-](https://www.ncei.noaa.gov/access/past-weather/) [weather/](https://www.ncei.noaa.gov/access/past-weather/)). Reported as the average of the daily minimum and maximum temperatures during the sampling period at each farm.

### Sample Collection

Blood samples were obtained weekly starting 6 wk before expected parturition, taken approximately at the time of feed delivery via puncture of coccygeal vessels using two-10mL evacuated tubes with serum separator (BD Vacutainer; Becton, Dickinson and Company, Franklin Lakes, NJ). Blood samples that were collected within 2 days of actual calving date were not considered as the -1 wk point to avoid changes in blood biomarkers due to the hormonal changes associated with calving, considering the previous week sample instead. Tubes were transported to the laboratory on ice, separated within 1 h by centrifugation at 2,000 × g for 20 min at 4°C, aliquoted, snap-frozen in liquid nitrogen, and stored at -80°C pending analysis within 1 month of collection.

Colostrum was harvested by trained farm personnel within 6 h of calving following each farm’s routine procedures. The volume of colostrum produced was measured using a graduated bucket (10 Quart Measuring Pail with Handle, United States Plastic Corporation, Lima, OH). A 50 mL sample of each cow’s colostrum was also collected and kept frozen at -20°C for further analysis.

### Analytical Determinations

#### Serum samples

Oxidant status was assessed following previously reported methods (Abuelo et al., 2016). The concentrations of reactive oxygen and nitrogen species (**RONS**) in serum were determined as indicator of pro-oxidant production using the OxiSelect In Vitro Reactive Oxygen and Nitrogen Species assay kit (Cell BioLabs Inc., San Diego, CA). Briefly, free radicals present in the sample bind to a dichlorodihydrofluorescein probe, converting it to a fluorescing product (2’,7’-dichlorodihydrofluorescein). Thus, the fluorescence intensity is proportional to the concentration of RONS in the sample. The fluorescence of dichlorofluorescent dye was determined at excitation wavelengths of 480 nm and emission of 530 nm in a Synergy H1 Hybrid plate reader (Biotek; Winooski, VT, USA). To ensure fluorescence at various concentrations, a standard curve, made by six serial dilutions (0–10,000 nM) of the fluorescence probe 2’,7’-dichlorodihydrofluorescein diacetate was included in each plate. All samples and standards were analyzed in duplicate and those with a CV greater than 10% were re-assayed. Background fluorescence was eliminated by subtracting blank values from sample values. Results are reported as the average relative fluorescence units (RFU) between replicates.

The antioxidant potential (**AOP**) of serum samples were determined using trolox (synthetic vitamin E analog) equivalents antioxidant capacity, as described previously (Re et al., 1999). Antioxidant components of serum interact, making it difficult to quantify each antioxidant individually. As a result, this method considers the synergism of all antioxidants present in a sample, including albumin, thiols, bilirubin, and superoxide dismutase. In brief, based on the standard curve of 0–25 g/L, a known volume of trolox standard concentration would result in a similar reduction of the radical 2,2’-azino-bis-3-ethylbenzothiazoline-6-sulfonic acid (Sigma-Aldrich, St. Louis, MO). Samples were analyzed in triplicate, and samples with replicates with CVs greater than 10% were re-assayed. Changes in oxidative balance may occur because of shifts in RONS and/or AOP. Thus, the oxidant status index (**OSi**) was calculated as the ratio between RONS and AOP, as this better characterizes shifts in redox balance in periparturient cows (Abuelo et al., 2013).

The serum concentration of BHB, blood urea nitrogen (**BUN)**, calcium (**Ca**), cholesterol (**Chol**), glucose (**Glu**), magnesium (**Mg**), nonesterified fatty acids (**NEFA**), albumin (**Alb**), and total protein (**TP**) were quantified using commercial reagents from Catachem Inc. (Bridgeport, CT) as biomarkers of nutrient utilization. As biomarker of inflammation, we determined the concentration of the positive acute phase protein Haptoglobin (**Hp**; Phase Haptoglobin Assay TP-801, Tridelta Development Limited, Maynooth, Ireland). All biomarkers related to nutrient utilization and inflammation were determined using a small-scale biochemistry analyzer (CataChemWell-T; Catachem Inc.) previously validated for cattle (Abuelo et al., 2020). The analyzer was calibrated every week using the assay manufacturer’s calibrators. Physiological and pathological reference samples were also analyzed at the time of calibration for two-level quality control. The precision of all biomarkers quantified in this analyzer is reported in Supplementary Table S3.

#### Colostrum samples

The concentration of IgG, IgA, and IgM in colostrum samples were measured via radial immunodiffusion (RID) (Bovine IgG, IgA and IgM test; Triple J Farms, Bellingham, WA) following manufacturer’s instructions (https://kentlabs.com/jjj/triple-j-farms-product-information/rid-plate-procedure/). The method is based on the precipitation in agarose gel growing in a circle antigen-antibody complexes which develop after 10-20 hours at room temperature and continues to grow until equilibrium is reached. Briefly, the colostrum samples were thawed overnight at 4 °C. Dilutions of each sample were performed in 0.9% NaCl. Samples were diluted at 1:6, 1:9, 1:10 for IgG analyses and at 1:2, 1:4, and 1:5 for IgA and IgM quantification. Standards were included in each plate for reference and ranged from 1.8 to 28.03 (IgG), 0.53 to 3.87 (IgA), and 0.62 to 3.81 (IgM) g/L. The diffusion ring through the agarose gel containing mono-specific antibody after 24 h of incubation at room temperature was measured using a caliper with a precision of 0.1 mm (VWR traceable caliper; Radnor, PA). The values of the sample’s ring were read off the standard curve to determine Ig concentrations in g/L. Samples falling outside of the standard curve were re-assayed using a higher or lower dilution as needed.

### Statistical analyses

Data were managed in Excel spreadsheets and exported to the statistical software. All statistical analyses were performed with JMP Pro v.15.2 (SAS Institute Inc., Cary, NC) and the criterion for statistical significance was established at *P* < 0.05. A two-way ANOVA was used to compare colostrum variables among seasons, lactation groups (first, second, or third to fifth), and farms. Associations between colostrum variables were examined using Spearman’s correlation coefficient. Cows were grouped *ex-post* into groups based on their colostrum variables to compare the changes in biomarkers of metabolic stress according to the volume and Ig content of their colostrum. Based on the recent recommendation of colostrum volume to sufficiently feed 2 meals of colostrum for one calf (Godden et al., 2019), we considered high colostrum producers (HCP) when cows produced ≥ 6 L while low colostrum producers (LCP) yielded < 6 L. Cows were classified as producing low (LIG) or high IgG (HIG) colostrum based on the industry threshold of 50 g/L (Godden, 2008). However, no industry standard exists for IgM and IgA to date. Thus, we used the median of the IgM and IgA values as the cutoff point to create equal sized groups of low and high Ig, LIM or HIM and LIA or HIA, respectively. Linear mixed models with repeated measures were built for the biomarkers Alb, BHBA, BUN, Ca, Chol, Glu, Hp, Mg, NEFA, TP, RONS, AOP and OSi as outcome variables. Fixed effects included time (sampling weeks -6 to -1 relative to actual calving), groups (LCP vs HCP, LIG vs HIG, LIM vs HIM, or LIA vs HIA) and their interaction. Cow nested within farm, season, and lactation group were used as random effects. For repeated measures, the covariance structures autoregressive 1, compound symmetry, or residual were tested for each variable, and the one with the lowest Akaike information criterion was chosen. Model assumptions were assessed by evaluation of homoscedasticity and normality of residuals. To satisfy these assumptions, some variables were natural log or square root-transformed and the resulting least squares means estimates were subsequently back transformed and presented as geometric means. All *P*-values given are those controlled for multiple comparisons with Tukey’s honestly significant difference test.

## RESULTS

### Descriptive results of colostrum variables

The distribution of colostrum volume and concentrations of IgG, IgM and IgA by season and lactation group is depicted in Figure 1. Distribution of cows per groups were 137 LCP vs. 91 HCP, 16 LIG vs. 212 HIG, 112 LIA vs. 116 HIA, and 113 LIM vs. 115 HIM. The overall average colostrum yield was 5.5 (range = 0.5 to 15.3) L with the highest average yield recorded in summer and lowest in winter (Table 2). Volume of colostrum did not vary among lactation groups (*P* = 0.24) or farms (*P* = 0.15). The overall colostrum average (range) IgG concentration was 118.7 (8.3 to 261.2) g/L, with cows calving in fall and winter producing more IgG concentrated colostrum than those calving in summer or spring (Table 2). Also, cows entering their 3^rd^ or greater lactation produced colostrum with greater IgG concentration than those entering their first or second lactation. No differences in IgG concentrations were found between first and second lactation animals. Colostrum IgG concentration was statistically different among all three farms of the study (*P* < 0.001).

**Figure 1:**
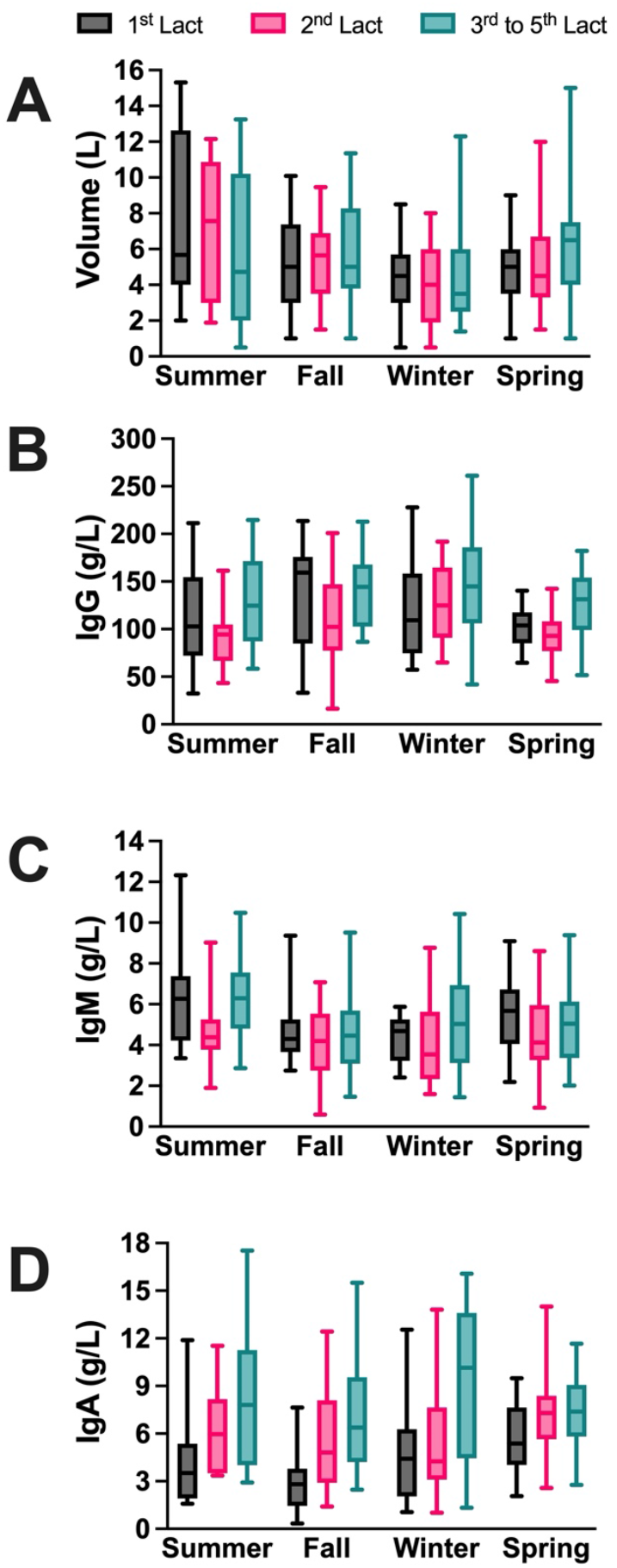
Distribution of colostrum (A) volume, (B) IgG concentration, (C) IgM concentration, and (D) IgA concentration per season and lactation group. The whiskers of the Box Plots represent the 5-95% percentiles.

**Table 2:** Distribution of colostrum variables across seasons, lactation groups, and study farms.

|  | Volume (L) | IgG (g/L) | IgM (g/L) | IgA (g/L) |
| --- | --- | --- | --- | --- |
| **Season** |  |  |  |  |
| *Summer* | 6.9 (6.6 – 7.3)<sup>a</sup> | 110.7 (105.5 – 115.9)<sup>a</sup> | 5.8 (5.5 – 6.0)<sup>a</sup> | 6.4 (6.0 – 6.8)<sup>a</sup> |
| *Fall* | 5.4 (5.1 – 5.8)<sup>b</sup> | 127.7 (122.6 – 132.8)<sup>b</sup> | 4.5 (4.2 – 4.7)<sup>b</sup> | 5.6 (5.2 – 6.0)<sup>b</sup> |
| *Winter* | 4.3 (4.0 – 4.6)<sup>c</sup> | 130.8 (125.9 – 135.6)<sup>b</sup> | 4.6 (4.4 – 4.8)<sup>bc</sup> | 6.4 (6.0 – 6.8)<sup>a</sup> |
| *Spring* | 5.4 (5.0 – 5.7)<sup>b</sup> | 104.9 (100.0 – 110.0)<sup>a</sup> | 5.0 (4.8 – 5.2)<sup>c</sup> | 6.8 (6.4 – 7.2)<sup>a</sup> |
| **Lactation group** |  |  |  |  |
| *1<sup>st</sup>* | 5.3 (5.0 – 5.6)<sup>a</sup> | 115.9 (111.4 – 120.4)<sup>a</sup> | 5.2 (5.0 – 5.5)<sup>a</sup> | 4.4 (4.0 – 4.8)<sup>a</sup> |
| *2<sup>nd</sup>* | 5.4 (5.1 – 5.7)<sup>a</sup> | 108.5 (104.8 – 112.2)<sup>a</sup> | 4.3 (4.1 – 4.5)<sup>b</sup> | 6.3 (6.0 – 6.6)<sup>b</sup> |
| *3<sup>rd</sup> to 5<sup>th</sup>* | 5.7 (5.4 – 6.0)<sup>a</sup> | 135.1 (130.8 – 139.4)<sup>b</sup> | 5.4 (5.2 – 5.6)<sup>a</sup> | 8.1 (7.8 – 8.4)<sup>c</sup> |
| **Farm** |  |  |  |  |
| *A* | 6.1 (5.7 – 6.9)<sup>a</sup> | 119.0 (114.7 – 123.4)<sup>a</sup> | 5.2 (5.0 – 5.4)<sup>a</sup> | 5.8 (5.5 – 6.2)<sup>a</sup> |
| *B* | 5.7 (5.4 – 6.0)<sup>a</sup> | 105.6 (101.0 – 110.1)<sup>b</sup> | 4.7 (4.5 – 4.9)<sup>b</sup> | 5.9 (5.5 – 6.2)<sup>a</sup> |
| *C* | 5.4 (5.1 – 5.8)<sup>a</sup> | 130.0 (125.8 – 134.2)<sup>c</sup> | 4.9 (4.7 – 5.1)<sup>ab</sup> | 7.1 (6.8 – 7.4)<sup>b</sup> |

2 Colostrum was harvested within 6 h of calving by farm staff. Results are expressed as means (95% confidence 3 intervals). Data were analyzed through two-way ANOVAs and means compared with Tukey honest significant 4 difference test. <sup>a,b,c</sup> Within each group and variable, means with different superscript letters are statistically 5 different ( $P < 0.05$ ).

The mean (range) IgM concentration of colostrum was 4.9 (0.6 to 12.3) g/L, with higher concentrations during summer (Table 2). Interestingly, cows entering the second lactation produced colostrum with significantly lower IgM concentration than those entering the first or third or greater (*P* < 0.023), and there were also differences in colostrum IgM concentration among farms (*P* = 0.019). For IgA, the overall mean (range) concentration was 6.29 (0.33 to 17.5) g/L, with lower concentrations recorded during fall compared to other months (Table 2). IgA concentration in colostrum increased with lactation number (*P* = 0.014), and Farm C’s colostrum had higher IgA concentrations than Farm A and B (*P* < 0.001). Colostrum volume showed a negative but weak correlation with the concentrations of IgG and IgA but did not correlate with IgM (Table 3). Conversely, the concentrations of all immunoglobulins showed a positive correlation with each other. Nevertheless, none of the correlations identified showed a correlation coefficient greater than 0.50.

**Table 3:** Correlations among colostrum volume and concentrations of IgG, IgM, and IgA.

| Variable | Volume (L) | IgG (g/L) | IgM (g/L) | IgA (g/L) |
| --- | --- | --- | --- | --- |
| Volume |  |  |  |  |
| $\rho$ | 1.0 | -0.28 | 0.03 | -0.15 |
| (95% CI) |  | (-0.38 to -0.15) | (-0.10 to 0.17) | (-0.28 to -0.02) |
| *P*-value |  | < 0.0001 | 0.62 | 0.021 |
| IgG |  |  |  |  |
| $\rho$ | | 1.0 | 0.26 | 0.50 |
| (95% CI) |  |  | (0.13 to 0.38) | (0.40 to 0.60) |
| *P*-value |  |  | < 0.0001 | < 0.0001 |
| IgM |  |  |  |  |
| $\rho$ | | | 1.0 | 0.30 |
| (95% CI) |  |  |  | (0.17 to 0.41) |
| *P*-value |  |  |  | < 0.0001 |
| IgA |  |  |  |  |
| $\rho$ | | | | 1.0 |
| (95% CI) |  |  |  |  |
| *P*-value |  |  |  |  |

7 Results expressed as Spearman's correlation coefficient ( $\rho$ ) with 95% confidence intervals (95% CI) and 8 associated *P*-values.

### Biomarkers of metabolic stress

The mean (SE) concentration of the cow biomarkers is presented by group and sampling point for the colostrum volume, IgG, IgM, and IgA variables in Tables 4, 5, 6 and 7, respectively. HCP cows had higher concentrations of AOP, BHB, and lower Chol and OSi than LCP. For IgG group, HIG cows showed higher concentrations of Alb and glucose compared to LIG. HIA cows had higher concentrations of Chol, RONS, OSi, and TP, whereas BHB and Glu were lower compared with LIA. There was also a tendency for lower NEFA concentrations (P = 0.096) in the HIA group. For the IgM group, biomarkers of the colostrum variables were not significantly different between HIM and LIM. Also, no significant differences were found on the biomarkers of inflammation haptoglobin and albumin for any of the colostrum variable categories analyzed.

**Table 4:** Estimated means and significance of the main effects from the generalized linear mixed models of blood biomarkers of late-gestation dairy cows comparing groups of high colostrum producers (HCP, n=91) and low colostrum producers (LCP, n=137).

| Outcome (units) | Estimated means of at the different time points relative to calving |  |  |  |  |  |  |  |  |  |  |  | SE | P value |  |  |
| --- | --- | --- | --- | --- | --- | --- | --- | --- | --- | --- | --- | --- | --- | --- | --- | --- |
|  | Wk -6 |  | Wk -5 |  | Wk -4 |  | Wk -3 |  | Wk -2 |  | Wk -1 |  |  | Group | Time | G×T |
|  | HCP | LCP | HCP | LCP | HCP | LCP | HCP | LCP | HCP | LCP | HCP | LCP |  |  |  |  |
| NEFA (mM)<sup>1</sup> | 0.22 | 0.24 | 0.21 | 0.21 | 0.20 | 0.22 | 0.19 | 0.22 | 0.19 | 0.21 | 0.23 | 0.23 | 0.08 | 0.491 | 0.027 | 0.555 |
| BHB (mg/dL)<sup>1</sup> | 4.23 | 4.09 | 4.39 | 4.05 | 4.32 | 4.03 | 4.51 | 3.90 | 4.65 | 3.78 | 5.15 | 4.17 | 0.30 | < 0.001 | < 0.001 | 0.001 |
| Cholesterol (mg/dL)<sup>1</sup> | 176 | 198 | 160 | 175 | 143 | 154 | 131 | 139 | 112 | 125 | 99 | 110 | 1.02 | 0.002 | < 0.001 | 0.214 |
| Glucose (mg/dL) | 73 | 76.6 | 74.3 | 74.8 | 72.6 | 73.0 | 72.8 | 73.3 | 72.0 | 74.5 | 71.7 | 73.2 | 1.01 | 0.222 | 0.003 | 0.087 |
| Total protein (g/dL)<sup>1</sup> | 7.54 | 7.41 | 7.62 | 7.37 | 7.45 | 7.24 | 7.40 | 7.23 | 7.14 | 7.10 | 6.97 | 6.82 | 0.29 | 0.153 | < 0.001 | 0.268 |
| BUN (mg/dL) | 11.4 | 10.8 | 11.8 | 11.00 | 11.72 | 11.22 | 11.46 | 11.85 | 11.99 | 12.32 | 12.94 | 12.86 | 0.37 | 0.646 | < 0.001 | 0.081 |
| Albumin (g/dL)<sup>1</sup> | 4.21 | 4.13 | 4.23 | 4.05 | 4.09 | 3.96 | 4.13 | 3.98 | 4.06 | 3.99 | 4.04 | 3.95 | 0.27 | 0.051 | < 0.001 | 0.343 |
| Haptoglobin (g/L)<sup>1</sup> | 0.50 | 0.46 | 0.49 | 0.48 | 0.47 | 0.45 | 0.50 | 0.43 | 0.48 | 0.44 | 0.46 | 0.42 | 0.05 | 0.229 | 0.019 | 0.205 |
| Calcium (mg/dL) | 9.29 | 9.16 | 9.27 | 9.18 | 9.03 | 9.02 | 9.28 | 9.06 | 9.10 | 9.17 | 9.21 | 9.11 | 0.11 | 0.533 | 0.016 | 0.213 |
| Magnesium (mg/dL) | 2.87 | 2.76 | 3.00 | 2.81 | 3.02 | 2.80 | 2.94 | 2.84 | 3.01 | 2.86 | 2.97 | 2.89 | 0.07 | 0.065 | 0.323 | 0.613 |
| AOP (TE/μL)<sup>1</sup> | 8.19 | 7.49 | 8.42 | 7.35 | 8.18 | 7.23 | 8.20 | 7.32 | 7.97 | 7.11 | 7.67 | 6.77 | 0.33 | 0.002 | < 0.001 | 0.690 |
| RONS (RFU)<sup>1</sup> | 44.8 | 49.3 | 46.8 | 49.8 | 48.2 | 51.5 | 52.6 | 55.3 | 54.9 | 59.2 | 53.7 | 61.7 | 1.05 | 0.239 | < 0.001 | 0.572 |
| OSi (arbitrary units)<sup>1</sup> | 5.38 | 6.53 | 5.46 | 6.73 | 5.79 | 7.07 | 6.31 | 7.50 | 6.78 | 8.27 | 6.88 | 9.05 | 0.06 | 0.006 | < 0.001 | 0.583 |

Generalized linear mixed models with repeated measurements were built for the blood biomarkers as outcomes with the Group, Time, their interaction (G×T) as fixed effects. Wk-6 through Wk -1 represent sample points relative to actual calving. HCP: High colostrum producers ( $\geq 6$ L); LCP: Low colostrum producers ( $< 6$ L). SE: Standard error. TE: Trolox equivalent; RFU: Relative fluorescence units; NEFA: non-esterified fatty acids; BHB: Beta-hydroxybutyrate; BUN: Blood urea nitrogen; AOP: Antioxidant potential; RONS: Reactive oxygen and nitrogen species; OSi: Oxidant status index. <sup>1</sup> Variable natural log transformed for statistical analysis.

**Table 5:** Estimated means and significance of the main effects from the generalized linear mixed models of blood biomarkers of late-gestation dairy cows comparing groups of high immunoglobulin G (HIG, n=212) and low immunoglobulin G (LIG, n=16).

| Outcome (units) | Estimated means of at the different time points relative to calving |  |  |  |  |  |  |  |  |  |  |  | SE | P value |  |  |
| --- | --- | --- | --- | --- | --- | --- | --- | --- | --- | --- | --- | --- | --- | --- | --- | --- |
|  | Wk -6 |  | Wk -5 |  | Wk -4 |  | Wk -3 |  | Wk -2 |  | Wk -1 |  |  | Group | Time | G×T |
|  | HIG | LIG | HIG | LIG | HIG | LIG | HIG | LIG | HIG | LIG | HIG | LIG |  |  |  |  |
| NEFA (mM)<sup>1</sup> | 0.23 | 0.13 | 0.22 | 0.15 | 0.21 | 0.15 | 0.20 | 0.18 | 0.20 | 0.18 | 0.23 | 0.24 | 0.09 | 0.304 | 0.328 | 0.415 |
| BHB (mg/dL)<sup>1</sup> | 4.12 | 3.75 | 4.19 | 3.52 | 4.13 | 4.19 | 4.13 | 3.88 | 4.11 | 3.90 | 4.51 | 5.08 | 0.71 | 0.605 | 0.004 | 0.249 |
| Cholesterol (mg/dL) | 196 | 198 | 176 | 154 | 154 | 141 | 139 | 129 | 124 | 110 | 109 | 93.3 | 8.4 | 0.369 | < 0.001 | 0.583 |
| Glucose (mg/dL)<sup>1</sup> | 75.5 | 68.7 | 74.9 | 66.9 | 73.4 | 64.7 | 73.4 | 69.6 | 73.9 | 66.4 | 72.9 | 67.4 | 1.11 | 0.013 | 0.464 | 0.548 |
| Total protein (g/dL)<sup>1</sup> | 7.48 | 7.20 | 7.51 | 7.02 | 7.35 | 6.98 | 7.32 | 7.13 | 7.14 | 6.76 | 6.91 | 6.51 | 0.18 | 0.165 | < 0.001 | 0.649 |
| BUN (mg/dL) | 11.0 | 12.1 | 11.2 | 12.8 | 11.3 | 13.7 | 11.5 | 13.8 | 12.1 | 13.6 | 12.8 | 13.8 | 1.14 | 0.089 | 0.101 | 0.766 |
| Albumin (g/dL)<sup>1</sup> | 4.17 | 3.84 | 4.13 | 3.85 | 4.02 | 3.80 | 4.04 | 3.88 | 4.02 | 3.86 | 4.00 | 3.58 | 0.02 | 0.043 | 0.018 | 0.172 |
| Haptoglobin (g/L)<sup>1</sup> | 0.48 | 0.56 | 0.48 | 0.45 | 0.46 | 0.45 | 0.46 | 0.54 | 0.45 | 0.43 | 0.44 | 0.45 | 0.10 | 0.816 | 0.242 | 0.367 |
| Calcium (mg/dL) | 9.23 | 8.92 | 9.22 | 9.09 | 9.04 | 8.60 | 9.14 | 9.05 | 9.15 | 8.87 | 9.15 | 9.00 | 0.21 | 0.430 | 0.263 | 0.820 |
| Magnesium (mg/dL)<sup>2</sup> | 2.77 | 2.87 | 2.86 | 2.92 | 2.86 | 2.96 | 2.85 | 2.86 | 2.88 | 3.04 | 2.91 | 2.57 | 0.15 | 0.941 | 0.270 | 0.122 |
| AOP (TE/μL)<sup>1</sup> | 7.85 | 7.09 | 7.84 | 7.21 | 7.67 | 6.82 | 7.75 | 6.85 | 7.52 | 6.73 | 7.21 | 6.18 | 0.36 | 0.213 | 0.001 | 0.914 |
| RONS (RFU)<sup>1</sup> | 47.7 | 34.1 | 48.7 | 41.4 | 50.5 | 37.8 | 54.3 | 47.2 | 57.5 | 50.3 | 58.2 | 57.2 | 2.11 | 0.282 | < 0.001 | 0.186 |
| OSi (arbitrary units)<sup>1</sup> | 6.01 | 4.81 | 6.15 | 5.73 | 6.51 | 5.53 | 6.94 | 6.89 | 7.57 | 7.47 | 8.00 | 9.26 | 1.13 | 0.771 | < 0.001 | 0.155 |

Generalized linear mixed models with repeated measurements were built for the blood biomarkers as outcomes with the Group, Time, their interaction (G×T) as fixed effects. Wk-6 through Wk -1 represent sample points relative to actual calving. HIG: High immunoglobulin G producers (≥ 50 g/L IgG); LIG: Low immunoglobulin G producers (< 50 g/L IgG). SE: Standard error. TE: Trolox equivalent; RFU: Relative fluorescence units; NEFA: non-esterified fatty acids; BHB: Beta-hydroxybutyrate; BUN: Blood urea nitrogen; AOP: Antioxidant potential; RONS: Reactive oxygen and nitrogen species; OSi: Oxidant status index. <sup>1</sup> Variable natural log transformed for statistical analysis. <sup>2</sup> Variable square root transformed for analysis.

**Table 6:** Estimated means and significance of the main effects from the generalized linear mixed models of blood biomarkers of late-gestation dairy cows comparing groups of high immunoglobulin M (HIM, n=115) and low immunoglobulin M (LIM, n=113).

| Outcome (units) | Estimated means of at the different time points relative to calving |  |  |  |  |  |  |  |  |  |  |  | SE | P value |  |  |
| --- | --- | --- | --- | --- | --- | --- | --- | --- | --- | --- | --- | --- | --- | --- | --- | --- |
|  | Wk -6 |  | Wk -5 |  | Wk -4 |  | Wk -3 |  | Wk -2 |  | Wk -1 |  |  | Group | Time | G×T |
|  | HIM | LIM | HIM | LIM | HIM | LIM | HIM | LIM | HIM | LIM | HIM | LIM |  |  |  |  |
| NEFA (mM)<sup>1</sup> | 0.23 | 0.23 | 0.23 | 0.20 | 0.21 | 0.21 | 0.21 | 0.20 | 0.19 | 0.21 | 0.23 | 0.23 | 0.08 | 0.925 | 0.040 | 0.477 |
| BHB (mg/dL)<sup>1</sup> | 4.01 | 4.23 | 4.11 | 4.22 | 4.20 | 4.09 | 4.27 | 4.00 | 4.17 | 4.07 | 4.47 | 4.60 | 0.29 | 0.966 | < 0.001 | 0.206 |
| Cholesterol (mg/dL)<sup>1</sup> | 187 | 188 | 167 | 170 | 150 | 147 | 135 | 135 | 118 | 121 | 105 | 105 | 2.15 | 0.885 | < 0.001 | 0.461 |
| Glucose (mg/dL)<sup>1</sup> | 75.3 | 74.8 | 74.3 | 74.5 | 73.1 | 72.6 | 72.2 | 73.9 | 73.1 | 73.6 | 71.8 | 73.2 | 1.01 | 0.709 | 0.004 | 0.437 |
| Total protein (g/dL)<sup>1</sup> | 7.73 | 7.50 | 7.50 | 7.45 | 7.35 | 7.3 | 7.32 | 7.29 | 7.13 | 7.10 | 6.94 | 6.83 | 0.59 | 0.752 | < 0.001 | 0.558 |
| BUN (mg/dL) | 11.0 | 11.1 | 11.4 | 11.1 | 11.8 | 10.9 | 12.1 | 11.2 | 12.4 | 12.0 | 12.9 | 12.8 | 0.36 | 0.315 | < 0.001 | 0.272 |
| Albumin (g/dL)<sup>1</sup> | 4.19 | 4.14 | 4.14 | 4.10 | 4.02 | 4.00 | 4.01 | 4.05 | 4.01 | 4.02 | 3.99 | 3.98 | 1.01 | 0.841 | < 0.001 | 0.643 |
| Haptoglobin (g/L)<sup>1</sup> | 0.46 | 0.51 | 0.48 | 0.48 | 0.45 | 0.47 | 0.48 | 0.45 | 0.45 | 0.45 | 0.43 | 0.45 | 0.23 | 0.714 | 0.007 | 0.098 |
| Calcium (mg/dL) | 9.12 | 9.31 | 9.13 | 9.29 | 8.97 | 9.08 | 9.06 | 9.22 | 9.08 | 9.20 | 9.08 | 9.20 | 0.41 | 0.251 | 0.025 | 0.992 |
| Magnesium (mg/dL) | 2.81 | 2.79 | 2.93 | 2.85 | 3.01 | 2.78 | 2.96 | 2.80 | 2.99 | 2.86 | 2.99 | 2.86 | 0.07 | 0.089 | 0.260 | 0.394 |
| AOP (TE/μL)<sup>1</sup> | 7.97 | 7.66 | 7.81 | 7.84 | 7.66 | 7.61 | 7.76 | 7.66 | 7.55 | 7.43 | 7.23 | 7.11 | 0.03 | 0.706 | < 0.001 | 0.775 |
| RONS (RFU)<sup>1</sup> | 50.0 | 44.0 | 51.7 | 45.1 | 53.4 | 46.6 | 56.8 | 51.2 | 57.1 | 56.9 | 60.6 | 55.6 | 1.45 | 0.156 | < 0.001 | 0.072 |
| OSi (arbitrary units)<sup>1</sup> | 6.19 | 5.71 | 6.53 | 5.73 | 6.88 | 6.08 | 7.23 | 6.65 | 7.47 | 7.61 | 8.28 | 7.77 | 0.06 | 0.323 | < 0.001 | 0.081 |

Generalized linear mixed models with repeated measurements were built for the blood biomarkers as outcomes with the Group, Time, their interaction (G×T) as fixed effects. Wk-6 through Wk -1 represent sample points relative to actual calving. HIM: High immunoglobulin M producers ( $\geq 4.6$ g/L IgG); LIM: Low immunoglobulin M producers ( $< 4.6$ g/L IgM). SE: Standard error. TE: Trolox equivalent; RFU: Relative fluorescence units; NEFA: non-esterified fatty acids; BHB: Beta-hydroxybutyrate; BUN: Blood urea nitrogen; AOP: Antioxidant potential; RONS: Reactive oxygen and nitrogen species; OSi: Oxidant status index. <sup>1</sup> Variable natural log transformed for statistical analysis.

**Table 7:** Estimated means and significance of the main effects from the generalized linear mixed models of blood biomarkers of late-gestation dairy cows comparing groups of high immunoglobulin A (HIA, n=116) and low immunoglobulin A (LIA, n=112).

| Outcome (units) | Estimated means of at the different time points relative to calving |  |  |  |  |  |  |  |  |  |  |  | SE | *P* value |  |  |
| --- | --- | --- | --- | --- | --- | --- | --- | --- | --- | --- | --- | --- | --- | --- | --- | --- |
|  | Wk -6 |  | Wk -5 |  | Wk -4 |  | Wk -3 |  | Wk -2 |  | Wk -1 |  |  | Group | Time | G×T |
|  | HIA | LIA | HIA | LIA | HIA | LIA | HIA | LIA | HIA | LIA | HIA | LIA |  |  |  |  |
| NEFA (mM)<sup>1</sup> | 0.23 | 0.24 | 0.19 | 0.25 | 0.19 | 0.23 | 0.18 | 0.23 | 0.18 | 0.22 | 0.23 | 0.23 | 0.08 | 0.096 | 0.030 | 0.046 |
| BHB (mg/dL)<sup>1</sup> | 3.89 | 4.29 | 4.02 | 4.25 | 3.99 | 4.23 | 4.15 | 4.05 | 3.73 | 4.47 | 4.12 | 4.93 | 0.03 | 0.007 | < 0.001 | < 0.001 |
| Cholesterol (mg/dL)<sup>1</sup> | 203 | 169 | 179 | 156 | 154 | 141 | 137 | 131 | 122 | 115 | 107 | 101 | 2.02 | 0.002 | < 0.001 | < 0.001 |
| Glucose (mg/dL)<sup>1</sup> | 73.8 | 76.3 | 73.1 | 75.7 | 71.0 | 74.8 | 70.8 | 75.2 | 72.0 | 74.8 | 70.8 | 74.2 | 1.13 | 0.005 | 0.003 | 0.663 |
| Total protein (g/dL)<sup>1</sup> | 7.66 | 7.24 | 7.69 | 7.22 | 7.46 | 7.11 | 7.39 | 7.13 | 7.29 | 6.86 | 7.01 | 6.66 | 0.09 | < 0.001 | < 0.001 | 0.090 |
| BUN (mg/dL) | 11.1 | 10.9 | 11.5 | 11.0 | 11.7 | 11.1 | 12.0 | 11.3 | 12.6 | 11.8 | 12.9 | 12.7 | 0.36 | 0.222 | < 0.001 | 0.682 |
| Albumin (g/dL)<sup>1</sup> | 4.18 | 4.14 | 4.13 | 4.11 | 4.01 | 4.01 | 3.99 | 4.07 | 4.04 | 3.99 | 4.02 | 3.95 | 0.01 | 0.782 | < 0.001 | 0.061 |
| Haptoglobin (g/L)<sup>1</sup> | 0.49 | 0.47 | 0.49 | 0.48 | 0.43 | 0.49 | 0.45 | 0.47 | 0.44 | 0.46 | 0.43 | 0.44 | 0.05 | 0.628 | 0.011 | 0.040 |
| Calcium (mg/dL) | 9.20 | 9.23 | 9.21 | 9.21 | 9.02 | 9.03 | 9.09 | 9.19 | 9.17 | 9.12 | 9.18 | 9.11 | 0.10 | 0.967 | 0.026 | 0.758 |
| Magnesium (mg/dL) | 2.77 | 2.79 | 2.82 | 2.92 | 2.84 | 2.89 | 2.84 | 2.87 | 2.88 | 2.91 | 2.83 | 2.96 | 0.02 | 0.409 | 0.189 | 0.747 |
| AOP (TE/μL)<sup>1</sup> | 7.94 | 7.67 | 7.71 | 7.92 | 7.79 | 7.47 | 7.75 | 7.65 | 7.59 | 7.37 | 7.24 | 7.08 | 0.05 | 0.6274 | < 0.001 | 0.083 |
| RONS (RFU)<sup>1</sup> | 54.2 | 40.8 | 56.0 | 41.8 | 58.4 | 42.6 | 62.4 | 47.0 | 65.3 | 50.1 | 66.7 | 50.9 | 1.05 | < 0.001 | < 0.001 | 0.928 |
| OSi (arbitrary units)<sup>1</sup> | 6.75 | 5.29 | 7.19 | 5.25 | 7.42 | 5.66 | 7.97 | 6.09 | 8.51 | 6.76 | 9.12 | 7.13 | 0.06 | < 0.001 | < 0.001 | 0.739 |

Generalized linear mixed models with repeated measurements were built for the blood biomarkers as outcomes with the Group, Time, their interaction (G×T) as fixed effects. Wk-6 through Wk -1 represent sample points relative to actual calving. HIA: High immunoglobulin A producers ( $\geq 5.52$ g/L IgA); LIG: Low immunoglobulin A producers ( $< 5.52$ g/L IgA). SE: Standard error. TE: Trolox equivalent; RFU: Relative fluorescence units; NEFA: non-esterified fatty acids; BHB: Beta-hydroxybutyrate; BUN: Blood urea nitrogen; AOP: Antioxidant potential; RONS: Reactive oxygen and nitrogen species; OSi: Oxidant status index. <sup>1</sup> Variable natural log transformed for statistical analysis.

## DISCUSSION

### Colostrum Variables

The averages and ranges in colostrum yield and Ig isotype concentrations found in our study are in line with previous reports (Kruse, 1970; Larson et al., 1980; Conneely et al., 2013; Quigley et al., 2013; Borchardt et al., 2022). Based on the group distributions, 93% of the cows in this study produced colostrum meeting the industry standard of IgG content (50 g/L), similar to the 96% documented by Conneely et al. (2013). Conversely, only 40% of cows produced sufficient first-milked colostrum to support a second meal of colostrum to calves (6 L total yield). Thus, suggesting that the volume of colostrum produced is a potential bottleneck for optimal colostrum feeding regimes in commercial dairy farms nowadays.

Studies reporting factors affecting colostrum volume are scarce in the literature. In agreement with two previous studies (Gavin et al., 2018; Borchardt et al., 2022), we found the lowest colostrum yield during winter. Nevertheless, contrary to previous reports (Conneely et al., 2013; Gavin et al., 2018), we did not note differences on colostrum yield by lactation group. A potential explanation for these differences is the broad range of time interval from calving to milking (0 to 21h) in the previous studies, compared to our study in which all cows were milked within the first 6 h after calving. It is known that colostrum IgG concentration decreases with time to harvest greater than 6-8 h postcalving (Conneely et al., 2013; Quigley et al., 2013). This is believed to be due to the dilution of colostrum due to the start of lactogenesis. Because parous cows produce more milk than primiparous cows, it is possible that this dilution effect is more marked in older cows, which could explain the differences in colostrum yield across parities observed in the Conneely et al. (2013) study as time of colostrum harvest is delayed. Pre-calving nutrition is also known to affect colostrum volume (Mann et al., 2016). Recently, Borchardt et al. (2022) also reported an association between duration in close-up diets and colostrum yield in a large dairy farm. Although we did not intend to evaluate nutrition factors influencing colostrum production, we purposedly selected farms with different dry-cow nutrition and management protocols, finding no differences in colostrum yield among study farms.

Overall, there was a marked individual variability in colostrum yield within seasons and lactation groups as noted by the broad confidence intervals in Figure 1. Thus, suggesting that many factors might influence colostrum yield production. Therefore, large multi-herd studies that investigate the epidemiology of colostrum production are urgently needed to identify which animals are more likely to produce sufficient amounts of high-quality colostrum.

The colostrum immunoglobulin concentration also showed important individual variability for all isotypes measured, but the ranges were similar to previous studies (Newby et al., 1982; Conneely et al., 2013). Immunoglobulin content varied by season. In agreement with previous reports, the lowest IgG concentration was documented in the spring (Conneely et al., 2013). The referenced study was conducted in a pasture-based dairy system and the authors speculated that the observed differences could be attributed to differences in dry period lengths or diet composition. However, we observed the same differences in IgG concentrations in farms that followed year-round calving patterns with a targeted dry period length and similar total mixed ration composition throughout the year. Thus, indicating that other underlying factors are likely involved, and further research is needed to elucidate the changes in IgG concentration associated with season of calving. Interestingly, during the fall and winter seasons, when colostrum showed the highest IgG concentrations, the concentrations of IgM were the lowest (Table 2) despite observing an overall positive correlation between IgG and IgM in colostrum samples (Table 3). Also, colostrum produced in the fall showed lower IgA content than at other seasons. To our knowledge, this is the first report evaluating changes in colostrum IgM and IgA concentrations across seasons and no research is available on factors affecting the colostrum concentration of these Ig isotypes.

Colostrum immunoglobulin composition also varied by parity. Cows entering their third to fifth lactation produced colostrum with greater IgG concentration compared to cows entering their first and second lactation. Interestingly, no differences were observed between parity 1 and 2 animals. Despite earlier studies recommending to discharge colostrum from first lactation heifers due to low IgG content (Selman et al., 1971), the results from this and previous studies do not support this (Conneely et al., 2013). The mean IgG concentration of colostrum for heifers in this study was twice that considered to be the threshold for good quality colostrum (50 g/L), and only 5% of the colostrum samples obtained from heifers were below that threshold. The higher IgG concentration in older (parity 3-5) cows is consistent with the existing literature (Kruse, 1970; Pritchett et al., 1991). Older cows are likely to have been exposed to a greater number of antigens in their lifetime, resulting in greater antibodies in serum and, subsequently, in colostrum (Larson et al., 1980). Although while IgG are transferred from the bloodstream across the mammary barrier into colostrum and IgA and IgM are largely derived from local synthesis by plasma cells in the mammary gland (Larson et al., 1980), differences in antigenic stimulation associated with age might also justify the observed increase in IgA concentration as parity increases. However, colostrum from parity 2 cows had lower IgM content than colostrum from parities 1 or 3-5. We cannot explain this finding based on the current evidence available of IgM content in colostrum. Colostrum Ig composition varied also by farm. However, these differences could be attributed to differences in management across farms (diet, housing, dry period length, etc.) as these factors are known to influence IgG content of colostrum (Godden, 2008).

Correlation analyses revealed a weak yet statistically significant negative association between colostrum volume and IgG (ρ = -0.28) and A (ρ = -0.15) concentrations. The negative association between colostrum yield and IgG concentration had also been previously reported (Quigley et al., 1994; Conneely et al., 2013; Mann et al., 2016). Nevertheless, the variation in colostrum volume only explained and 7.8 and 2.3% of the variation in colostrum IgG and IgA concentrations, respectively (Table 3). Thus, indicating a weak association between colostrum volume and immunoglobulin concentration. This suggests that the processes of synthesis and transfer of IgG and IgA to colostrum might be largely independent of the volume of colostrum produced. For example, colostral IgG are derived from those circulating in plasma and are actively taken up by the mammary gland through binding to FcRn receptors (Zhang et al., 2009); whereas the volume of produced milk depends on the osmotic equilibrium of the blood–milk barrier, regulated mainly by lactose (Costa et al., 2019). We speculate that the amount of absorbed water in the mammary gland alveoli during colostrogenesis, and thus, the colostrum volume is also dependent on this equilibrium, which is affected my many other solutes beyond Igs.

Traditionally, colostrum quality has been evaluated based solely on the concentration of IgG as this is the most abundant immunoglobulin isotype in bovine colostrum (Larson et al., 1980). However, we found weak correlations (defined as Spearman’s coefficient ≤ 0.50) among the different isotypes of immunoglobulins in colostrum. Thus, optimization of passive immunity transfer for IgG might not necessarily result in optimal transfer of other immunological factors of colostrum, such as other Ig isotypes. However, the relevance of the transfer of IgA and IgM to the calf via colostrum on calf health and productivity remains unexplored to date and more research is needed to unravel the impact that all colostrum immunological components have on calf health.

### Nutrient Utilization and Colostrum Variables

We evaluated the association between colostrum variables and biomarkers of energy (NEFA, BHB, Glu, Chol), protein (BUN, TP), and macromineral (Ca, Mg) status in blood samples collected throughout the last 6 wk of gestation. The higher BHB concentration exhibited by HCP cows compared to LCP indicates that cows producing > 6 L of colostrum had greater nutrient demands than those producing less than 6 L, as BHB is commonly used as an indicator of energy deficit (McArt et al., 2013). However, the lack of differences in NEFA concentration between colostrum volume groups suggests that cows were able to cope with these increased energy demands without a marked increase in lipid mobilization. Furthermore, HCP cows exhibited lower Chol concentrations than LCP cows, which can be associated with decreased hepatic Chol export due to increased ketogenesis (Kessler et al., 2014; Gross et al., 2021), and supports the finding of increased energy demands in association with the volume of colostrum produced, despite no marked lipid mobilization.

Changes in nutrient utilization biomarkers were also identified between the colostrum IgG and IgA groups but not for IgM. Colostral IgG are derived from those circulating in plasma and are taken up by the mammary gland via action of receptors (Zhang et al., 2009). Changes in colostrum IgG concentrations could therefore be caused by changes in circulating blood concentrations in the dam, a change in transfer capability, a difference in the rate of water inclusion, or a combination of these. Cows producing colostrum above the 50 g/L IgG cut-off showed greater Glu concentrations than those producing colostrum below this threshold, but no other differences in energy-related metabolites were identified. In a recent study, Immler et al. (2021) also found no relationship the Brix value of the colostrum (an estimate of IgG concentration) and the biomarkers Chol or NEFA. However, these authors did not include Glu in their panel of biomarkers. Glu is the primary fuel for immune cells (Calder et al., 2007), and therefore it is plausible that greater availability of glucose as energy source allowed blood lymphocytes to synthetize more IgG that would have been subsequently transported into the mammary gland for colostrum synthesis. In fact, a positive correlation between plasma glucose and IgG has been documented in goats (Hefnawy et al., 2010). However, studies in human and bovine lymphocytes described a reduction in lymphocyte function under high concentrations of glucose in vitro (Franklin et al., 1991; Jennbacken et al., 2013). However, these studies used concentrations of glucose (11.1 m*M*) exceeding normal blood concentrations in cattle (∼ 3.2 – 4.4 m*M*), limiting the translation of their findings to the live animal. To our knowledge, the biological significance of blood glucose concentrations on immunoglobulin synthesis has not been investigated in cattle to date. Furthermore, we did not assess blood IgG concentrations in the cows in the present study, which limits our ability to discern if the changes in IgG colostrum content is associated with differences in blood IgG concentrations or due to other steps involved in the translocation if IgG from bloodstream into colostrum.

Cows with higher IgA colostrum concentration showed a metabolic profile suggestive of a better energy status (lower concentrations of BHB and a tendency towards lower NEFA, and higher concentrations of Chol and TP). Unlike IgG, IgA are primarily synthetized locally in the mammary gland and not translocated from bloodstream (Larson et al., 1980). However, it is possible that the greater energy balance would allow B lymphocytes in the mammary gland to secrete more IgA, as producing IgA is a considerable energy expense for the body of mammals (Woof and Kerr, 2006). However, it could also be possible that the greater IgA concentration in colostrum was due to increased recruitment of B cells into the mammary gland and/or increased class switch to IgA^+^ cells. Lower serum Glu concentrations were also found in cows with higher IgA colostrum. Because the mammary gland uptakes 60-85% of blood glucose (Annison and Linzell, 1964; Rigout et al., 2002), lower glycemia might be indicative of higher availability of glucose for the mammary gland IgA-producing immune cells, as Glu is the main fuel used by these cells (Calder et al., 2007). We are unaware of any research in the bovine species studying the impact of energy status on the colostrogenesis mechanisms, and, therefore, more research is needed in this area to be able to modulate colostrum production in the dairy cow.

We detected no association between any of the colostrum variables studied and concentrations of the macrominerals Ca or Mg. Immler et al. (2021), however, found a negative association between serum Ca concentration and colostrum Brix value was, but were unable to clarify the physiological background behind this observation. Thus, the differences between their study and ours underscores the need for further research in this area, given that different Ca management nutritional strategies resulted in differences in colostrum IgG concentrations (Diehl et al., 2018).

### Inflammatory Status and Colostrum Variables

Based on the biomarkers Hp and Alb, positive and negative acute phase proteins respectively, we observed no differences in the inflammatory status of cows based on colostrum volume or Ig content. This suggests that the cows’ physiological homeostasis was not disrupted during late gestation in association with the colostrum variables studied. Although exacerbated inflammation is often seen in transition cows (Bradford et al., 2015), and is one of the hallmarks of metabolic stress (Abuelo et al., 2019), changes in biomarkers of inflammation usually occur after parturition (Abuelo et al., 2014; Burfeind et al., 2014; Pohl et al., 2015). Thus, even though we are unable to determine cause-effect relationships in this observational study, the lack of association between acute phase proteins and colostrum yield or Ig isotype content let us to speculate that the colostrogenesis process might not increase inflammation in the pre-partum cow.

### Oxidant Status and Colostrum Variables

Oxidative stress is also a common feature of the transition period of dairy cattle (Abuelo et al., 2015), known to impair functional capabilities of immune cell populations, including the lymphocytes responsible for Ig synthesis (Lacetera et al., 2005; Sordillo and Aitken, 2009; Cuervo et al., 2021). Thus, we anticipated that lower colostrum Ig content or volume would be associated with a more pro-oxidant systemic redox balance. In line with our hypothesis, LCP cows showed greater OSi values than HCP cows. This was due to differences in AOP as RONS remained similar between colostrum yield groups (Table 4). Therefore, it is possible that higher AOP might have supported HCP cows to produce more colostrum, rendering availability of antioxidants a potential limiting factor in colostrum volume production for cows. Our study is, to our knowledge, the first one to examine relationships between cow oxidant status and colostrum production, but given the observed relationship between lower AOP and colostrum volume, the extent to which increasing antioxidant capacity in late-gestation cows enhances colostrum yield warrants further research.

There were no differences in oxidant status between cows in the IgG or IgM groups. However, HIA cows showed higher concentration of RONS and greater OSi values compared to LIA. This finding is contrary to our initial hypothesis linking a pro-oxidant status to lower Ig content. How systemic redox balance might influence the production of the mucosa-derived IgA but not IgG or IgM remains unexplained and asserts the complexity of redox regulation of biological processes. Certainly, the role of oxidative balance in the colostrogenesis process needs to be elucidated further.

### Study Limitations

Given the observational nature of this study, we were only able to show associations among colostrum variables and biomarkers of metabolic stress and did not explore cause-effect relationships. However, this is the first study to investigate changes in biomarkers of metabolic stress in association with colostrum variables and the associations identified can be examined further via controlled interventional studies. Another limitation of the study is that using the industry standard of colostrum IgG concentration (50 g/L) resulted in unbalanced groups sizes (212 vs. 16 cows in HIG and LIG), which might have influenced our ability to detect differences. In fact, post hoc power calculations revealed that only powers of 0.13, 0.21, 0.11, and 0.19 for detecting differences in NEFA concentrations between groups of colostrum volume, IgG, IgM, and IgA, respectively. Thus, we cannot exclude associations between metabolic stress and IgG content that we were identified in this study.

## CONCLUSIONS

This study evaluated, for the first time, the metabolic status of dairy cows during the last 6 wk of gestation based on the volume and Ig concentration of colostrum. We observed marked individual variability in colostrum yield and Ig isotype concentration, suggesting that colostrum production is a complex and multifactorial process. Also, we detected differences in nutrient utilization and oxidant status biomarkers in association with the volume and IgG and IgA concentration of colostrum. Among all changes detected, our results suggest that increasing availability of antioxidants during late-gestation could support the production of higher volumes of colostrum, which warrants further investigation through supplementation trials.

## ACKNOWLEDGEMENTS

This research was funded by the USDA National Institute of Food and Agriculture Animal Health project number 1016161 and a grant from the Michigan Alliance for Animal Agriculture. The funders had no role in the design and conduct of the study; collection, management, analysis, and interpretation of the data; preparation, review, or approval of the manuscript; and decision to submit the manuscript for publication. The authors declare no conflicts of interest.

## REFERENCES

Abuelo, A., J. Hernandez, J. L. Benedito, and C. Castillo. 2013. Oxidative stress index (OSi) as a new tool to assess redox status in dairy cattle during the transition period. Animal 7(8):1374–1378. http://doi.org/10.1017/S1751731113000396

Abuelo, A., J. Hernandez, J. L. Benedito, and C. Castillo. 2014. A comparative study of the metabolic profile, insulin sensitivity and inflammatory response between organically and conventionally managed dairy cattle during the periparturient period. Animal 8(9):1516–1525. http://doi.org/10.1017/S1751731114001311

Abuelo, A., J. Hernandez, J. L. Benedito, and C. Castillo. 2015. The importance of the oxidative status of dairy cattle in the periparturient period: revisiting antioxidant supplementation. J. Anim. Physiol. Anim. Nutr. (Berl) 99(6):1003–1016. http://doi.org/10.1111/jpn.12273

Abuelo, A., J. C. Gandy, L. Neuder, J. Brester, and L. M. Sordillo. 2016. Short communication: Markers of oxidant status and inflammation relative to the development of claw lesions associated with lameness in early lactation cows. J. Dairy Sci. 99(7):5640–5648. http://doi.org/10.3168/jds.2015-10707

Abuelo, A., J. Hernandez, J. L. Benedito, and C. Castillo. 2019. Redox Biology in Transition Periods of Dairy Cattle: Role in the Health of Periparturient and Neonatal Animals. Antioxidants (Basel) 8(1):20. http://doi.org/10.3390/antiox8010020

Abuelo, A., J. L. Brester, K. Starken, and L. M. Neuder. 2020. Technical note: Comparative evaluation of 3 methods for the quantification of nonesterified fatty acids in bovine plasma sampled prepartum. J. Dairy Sci. 103(3):2711–2717. http://doi.org/10.3168/jds.2019-17527

Abuelo, A., F. Cullens, A. Hanes, and J. L. Brester. 2021. Impact of 2 Versus 1 Colostrum Meals on Failure of Transfer of Passive Immunity, Pre-Weaning Morbidity and Mortality, and Performance of Dairy Calves in a Large Dairy Herd. Animals (Basel) 11(3):782. http://doi.org/10.3390/ani11030782

Annison, E. F. and J. L. Linzell. 1964. The Oxidation and Utilization of Glucose and Acetate by the Mammary Gland of the Goat in Relation to Their over-All Metabolism and Milk Formation. J Physiol 175(3):372–385. http://doi.org/10.1113/jphysiol.1964.sp007522

Barrington, G. M., T. B. McFadden, M. T. Huyler, and T. E. Besser. 2001. Regulation of colostrogenesis in cattle. Livestock Production Science 70(1-2):95–104. http://doi.org/10.1016/s0301-6226(01)00201-9

Baumrucker, C. R. and R. M. Bruckmaier. 2014. Colostrogenesis: IgG1 transcytosis mechanisms. J Mammary Gland Biol Neoplasia 19(1):103–117. http://doi.org/10.1007/s10911-013-9313-5

Borchardt, S., F. Sutter, W. Heuwieser, and P. Venjakob. 2022. Management-related factors in dry cows and their associations with colostrum quantity and quality on a large commercial dairy farm. J. Dairy Sci. 105(2):1589–1602. http://doi.org/10.3168/jds.2021-20671

Bradford, B. J., K. Yuan, J. K. Farney, L. K. Mamedova, and A. J. Carpenter. 2015. Invited review: Inflammation during the transition to lactation: New adventures with an old flame. J. Dairy Sci. 98(10):6631–6650. http://doi.org/10.3168/jds.2015-9683

Burfeind, O., I. Sannmann, R. Voigtsberger, and W. Heuwieser. 2014. Receiver operating characteristic curve analysis to determine the diagnostic performance of serum haptoglobin concentration for the diagnosis of acute puerperal metritis in dairy cows. Anim Reprod Sci 149(3-4):145–151. http://doi.org/10.1016/j.anireprosci.2014.07.020

Calder, P. C., G. Dimitriadis, and P. Newsholme. 2007. Glucose metabolism in lymphoid and inflammatory cells and tissues. Current Opinion in Clinical Nutrition and Metabolic Care 10(4):531–540. http://doi.org/10.1097/MCO.0b013e3281e72ad4

Conneely, M., D. P. Berry, R. Sayers, J. P. Murphy, I. Lorenz, M. L. Doherty, and E. Kennedy. 2013. Factors associated with the concentration of immunoglobulin G in the colostrum of dairy cows. Animal 7(11):1824–1832. http://doi.org/10.1017/S1751731113001444

Costa, A., N. Lopez-Villalobos, N. W. Sneddon, L. Shalloo, M. Franzoi, M. De Marchi, and M. Penasa. 2019. Invited review: Milk lactose—Current status and future challenges in dairy cattle. J. Dairy Sci. 102(7):5883–5898. http://doi.org/https://doi.org/10.3168/jds.2018-15955

Cuervo, W., L. M. Sordillo, and A. Abuelo. 2021. Oxidative Stress Compromises Lymphocyte Function in Neonatal Dairy Calves. Antioxidants (Basel) 10(2):255. http://doi.org/10.3390/antiox10020255

Diehl, A. L., J. K. Bernard, S. Tao, T. N. Smith, T. Marins, D. J. Kirk, D. J. McLean, and J. D. Chapman. 2018. Short communication: Blood mineral and gas concentrations of calves born to cows fed prepartum diets differing in dietary cation-anion difference and calcium concentration. J. Dairy Sci. 101(10):9048–9051. http://doi.org/10.3168/jds.2018-14829

Franklin, S. T., J. W. Young, and B. J. Nonnecke. 1991. Effects of Ketones, Acetate, Butyrate, and Glucose on Bovine Lymphocyte Proliferation1,2. J. Dairy Sci. 74(8):2507–2514. http://doi.org/https://doi.org/10.3168/jds.S0022-0302(91)78428-2

Gavin, K., H. Neibergs, A. Hoffman, J. N. Kiser, M. A. Cornmesser, S. A. Haredasht, B. Martinez-Lopez, J. R. Wenz, and D. A. Moore. 2018. Low colostrum yield in Jersey cattle and potential risk factors. J. Dairy Sci. 101(7):6388–6398. http://doi.org/10.3168/jds.2017-14308

Godden, S. 2008. Colostrum management for dairy calves. Vet. Clin. North. Am. Food Anim. Pract. 24(1):19–39. http://doi.org/10.1016/j.cvfa.2007.10.005

Godden, S. M., J. E. Lombard, and A. R. Woolums. 2019. Colostrum Management for Dairy Calves. Vet. Clin. North. Am. Food Anim. Pract. 35(3):535–556. http://doi.org/10.1016/j.cvfa.2019.07.005

Gross, J. J., A. C. Schwinn, E. Müller, A. Münger, F. Dohme-Meier, and R. M. Bruckmaier. 2021. Plasma cholesterol levels and short-term adaptations of metabolism and milk production during feed restriction in early lactating dairy cows on pasture. J. Anim. Physiol. Anim. Nutr. (Berl) 105(6):1024–1033. http://doi.org/10.1111/jpn.13531

Hammon, H. M., J. Steinhoff-Wagner, J. Flor, U. Schonhusen, and C. C. Metges. 2013. Lactation Biology Symposium: role of colostrum and colostrum components on glucose metabolism in neonatal calves. J. Anim. Sci. 91(2):685–695. http://doi.org/10.2527/jas.2012-5758

Hefnawy, A.-E., S. Youssef, and S. Shousha. 2010. Some Immunohormonal Changes in Experimentally Pregnant Toxemic Goats. Veterinary Medicine International 2010:768438. http://doi.org/10.4061/2010/768438

Immler, M., K. Failing, T. Gartner, A. Wehrend, and K. Donat. 2021. Associations between the metabolic status of the cow and colostrum quality as determined by Brix refractometry. J. Dairy Sci. 104(9):10131–10142. http://doi.org/10.3168/jds.2020-19812

Jennbacken, K., S. Ståhlman, L. Grahnemo, O. Wiklund, and L. Fogelstrand. 2013. Glucose impairs B-1 cell function in diabetes. Clinical and Experimental Immunology 174(1):129–138. http://doi.org/10.1111/cei.12148

Kehrli Jr, M. E., B. J. Nonnecke, and J. A. Roth. 1989. Alterations in bovine neutrophil function during the periparturient period. Am. J. Vet. Res. 50(2):207–214. https://www.scopus.com/inward/record.uri?eid=2-s2.0-0024617208&partnerID=40&md5=29e7b634c193b280a19f03237d918533

Kessler, E. C., J. J. Gross, R. M. Bruckmaier, and C. Albrecht. 2014. Cholesterol metabolism, transport, and hepatic regulation in dairy cows during transition and early lactation. J. Dairy Sci. 97(9):5481–5490. http://doi.org/https://doi.org/10.3168/jds.2014-7926

Kessler, E. C., G. C. Pistol, R. M. Bruckmaier, and J. J. Gross. 2020. Pattern of milk yield and immunoglobulin concentration and factors associated with colostrum quality at the quarter level in dairy cows after parturition. J. Dairy Sci. 103(1):965–971. http://doi.org/10.3168/jds.2019-17283

Kruse, V. 1970. Yield of colostrum and immunoglobulin in cattle at the first milking after parturition. Animal Science 12(4):619–626. http://doi.org/10.1017/S0003356100029263

Lacetera, N., D. Scalia, U. Bernabucci, B. Ronchi, D. Pirazzi, and A. Nardone. 2005. Lymphocyte functions in overconditioned cows around parturition. J. Dairy Sci. 88(6):2010–2016. http://doi.org/10.3168/jds.S0022-0302(05)72877-0

Larson, B. L., H. L. Heary, Jr., and J. E. Devery. 1980. Immunoglobulin production and transport by the mammary gland. J. Dairy Sci. 63(4):665–671. http://doi.org/10.3168/jds.S0022-0302(80)82988-2

Mann, S., F. A. Leal Yepes, T. R. Overton, A. L. Lock, S. V. Lamb, J. J. Wakshlag, and D. V. Nydam. 2016. Effect of dry period dietary energy level in dairy cattle on volume, concentrations of immunoglobulin G, insulin, and fatty acid composition of colostrum. J. Dairy Sci. 99(2):1515–1526. http://doi.org/10.3168/jds.2015-9926

McArt, J. A., D. V. Nydam, G. R. Oetzel, T. R. Overton, and P. A. Ospina. 2013. Elevated non-esterified fatty acids and beta-hydroxybutyrate and their association with transition dairy cow performance. Vet J 198(3):560–570. http://doi.org/10.1016/j.tvjl.2013.08.011

Morin, D. E., G. C. McCoy, and W. L. Hurley. 1997. Effects of quality, quantity, and timing of colostrum feeding and addition of a dried colostrum supplement on immunoglobulin G1 absorption in Holstein bull calves. J. Dairy Sci. 80(4):747–753. http://doi.org/10.3168/jds.S0022-0302(97)75994-0

Newby, T. J., C. R. Stokes, and F. J. Bourne. 1982. Immunological activities of milk. Vet. Immunol. Immunopathol. 3(1-2):67–94. http://doi.org/10.1016/0165-2427(82)90032-0

NRC, National Research Council. 2001. Nutrient requirements of dairy cattle. 7th ed. National Academic Press, Washington DC, USA.

Pohl, A., O. Burfeind, and W. Heuwieser. 2015. The associations between postpartum serum haptoglobin concentration and metabolic status, calving difficulties, retained fetal membranes, and metritis. J. Dairy Sci. 98(7):4544–4551. http://doi.org/10.3168/jds.2014-9181

Pritchett, L. C., C. C. Gay, T. E. Besser, and D. D. Hancock. 1991. Management and production factors influencing immunoglobulin G1 concentration in colostrum from Holstein cows. J. Dairy Sci. 74(7):2336–2341. http://doi.org/10.3168/jds.S0022-0302(91)78406-3

Quigley, J. D., K. R. Martin, H. H. Dowlen, L. B. Wallis, and K. Lamar. 1994. Immunoglobulin Concentration, Specific Gravity, and Nitrogen Fractions of Colostrum from Jersey Cattle1. J. Dairy Sci. 77(1):264–269. http://doi.org/https://doi.org/10.3168/jds.S0022-0302(94)76950-2

Quigley, J. D., A. Lago, C. Chapman, P. Erickson, and J. Polo. 2013. Evaluation of the Brix refractometer to estimate immunoglobulin G concentration in bovine colostrum. J. Dairy Sci. 96(2):1148–1155. http://doi.org/10.3168/jds.2012-5823

Raboisson, D., P. Trillat, and C. Cahuzac. 2016. Failure of Passive Immune Transfer in Calves: A Meta-Analysis on the Consequences and Assessment of the Economic Impact. PLoS One 11(3):e0150452. http://doi.org/10.1371/journal.pone.0150452

Re, R., N. Pellegrini, A. Proteggente, A. Pannala, M. Yang, and C. Rice-Evans. 1999. Antioxidant activity applying an improved ABTS radical cation decolorization assay. Free Radic. Biol. Med. 26(9-10):1231–1237. http://doi.org/10.1016/s0891-5849(98)00315-3

Rigout, S., S. Lemosquet, J. E. van Eys, J. W. Blum, and H. Rulquin. 2002. Duodenal Glucose Increases Glucose Fluxes and Lactose Synthesis in Grass Silage-Fed Dairy Cows. J. Dairy Sci. 85(3):595–606. http://doi.org/https://doi.org/10.3168/jds.S0022-0302(02)74113-1

Selman, I. E., A. D. McEwan, and E. W. Fisher. 1971. Absorption of immune lactoglobulin by newborn dairy calves. Attempts to produce consistent immune lactoglobulin absorptions in newborn dairy calves using standardised methods of colostrum feeding and management. Res Vet Sci 12(3):205–210.

Sordillo, L. M. and S. L. Aitken. 2009. Impact of oxidative stress on the health and immune function of dairy cattle. Vet. Immunol. Immunopathol. 128(1-3):104–109. http://doi.org/10.1016/j.vetimm.2008.10.305

Sordillo, L. M. and V. Mavangira. 2014. The nexus between nutrient metabolism, oxidative stress and inflammation in transition cows. Anim. Prod. Sci. 54(9):1204–1214. http://doi.org/10.1071/an14503

Urie, N. J., J. E. Lombard, C. B. Shivley, C. A. Kopral, A. E. Adams, T. J. Earleywine, J. D. Olson, and F. B. Garry. 2018. Preweaned heifer management on US dairy operations: Part V. Factors associated with morbidity and mortality in preweaned dairy heifer calves. J. Dairy Sci. 101(10):9229–9244. http://doi.org/10.3168/jds.2017-14019

Wildman, E. E., G. M. Jones, P. E. Wagner, R. L. Boman, H. F. Troutt, and T. N. Lesch. 1982. A Dairy-Cow Body Condition Scoring System and Its Relationship to Selected Production Characteristics. J. Dairy Sci. 65(3):495–501. http://doi.org/DOI 10.3168/jds.S0022-0302(82)82223-6

Woof, J. M. and M. A. Kerr. 2006. The function of immunoglobulin A in immunity. J Pathol 208(2):270–282. http://doi.org/10.1002/path.1877

Zhang, R., Z. Zhao, Y. Zhao, I. Kacskovics, M. Eijk, N. Groot, N. Li, and L. Hammarstrom. 2009. Association of FcRn Heavy Chain Encoding Gene (FCGRT) Polymorphisms with IgG Content in Bovine Colostrum. Anim Biotechnol 20(4):242–246. http://doi.org/10.1080/10495390903196448

