## Supplementary material for "Changes in biomarkers of metabolic stress during late gestation of dairy cows associated with colostrum volume and immunoglobulin content": Table S1

### SUPPLEMENTARY MATERIALS

**Table S1: Ingredient and nutrient composition of diets as formulated.**

| | Prepartum diet composition | | | | | | | |
|---|---|---|---|---|---|---|---|---|
| | Farm A | | | | Farm B | | | Farm C |
| | Close up cow | Close up heifer | Far off cow | Far off heifer | Close up | Far off cow | Far off heifer | Dry cow / heifer |
| **Total DM, kg** | 13.15 | 11.3 | 12.5 | 11.3 | 12.9 | 13.0 | 10.6 | 12.7 / 10.9 |
| **Ingredient, % of DM** | | | | | | | | |
| Corn Silage | 34.3 | 40.1 | 18.2 | 18.2 | 53.47 | 20.91 | - | 40.5 |
| Sorghum Silage | - | - | - | - | - | 8.71 | 30 | - |
| Beet Pulp Wet | 12.9 | - | - | - | - | - | - | - |
| Haylage | - | - | 69.2 | 69.2 | - | 48.79 | 54.44 | 12.8 |
| Wheat Straw | - | 28 | 10.9 | 10.9 | 20.58 | 19.17 | - | 37.6 |
| Grass hay | 21.6 | - | - | - | - | - | - | - |
| Lact TMR Leftover | - | - | - | - | - | - | 12.34 | - |
| Mineral premix | 31.2 | 31.9 | 1.6 | 1.6 | 25.95 | 2.07 | 3.21 | 9.1 |
| Salt | - | - | - | - | - | 0.35 | - | - |
| **Nutrient composition, DM basis** | | | | | | | | |
| CP, % | 14.2 | 15.9 | 13.8 | 13.8 | 14.4 | 13.1 | 16.3 | 12.3 |
| MP, % | 3.1 | 4.2 | 1.5 | 1.5 | 7.9 | 5.8 | 7.6 | 5.3 |
| Soluble Protein, % | 4.8 | 6.3 | 7.8 | 7.8 | 6.2 | 3.4 | 4.1 | 6.1 |
| Lys/Met, - | 3.4 | 3.8 | 3.5 | 3.5 | 4.7 | 5.6 | 4.3 | 4.1 |
| RUP, % | 4.1 | 5.6 | 3.6 | 3.6 | 5.0 | 3.3 | 4.4 | 3.8 |
| ME, Mcal/kg | 2.32 | 2.34 | 2.05 | 2.05 | 2.38 | 2.23 | 2.43 | 2.16 |
| ADF, % | 23.7 | 28.3 | 40.9 | 40.9 | 24.4 | 36.6 | 34.6 | 35.8 |
| NDF, % | 39.0 | 44.7 | 49.2 | 49.2 | 40.1 | 49.3 | 44.0 | 48.3 |
| Forage NDF, % | 26.8 | 36.7 | 48.9 | 48.9 | 35.8 | 48.9 | 42.0 | 45.7 |
| Starch, % | 18.6 | 18.5 | 7.3 | 7.3 | 23.3 | 9.5 | 5.0 | 9.3 |
| Sugar, % | 4.1 | 1.8 | 3.4 | 3.4 | 2.3 | 3.3 | 5.4 | 5.1 |
| Fat, % | 2.5 | 2.7 | 2.6 | 2.6 | 2.5 | 3.0 | 3.6 | 2.8 |
| PUFA, % | 0.81 | 0.92 | 0.55 | 0.55 | 0.96 | 0.78 | 0.83 | 0.73 |
| Ca, % | 1.7 | 0.8 | 1.0 | 1.0 | 1.2 | 0.8 | 1.0 | 0.8 |

| | | | | | | | | |
|---|---|---|---|---|---|---|---|---|
| P, % | 0.42 | 0.35 | 0.28 | 0.28 | 0.38 | 0.32 | 0.36 | 0.34 |
| Ca:P ratio, - | 4.1 | 2.4 | 3.6 | 3.6 | 3.2 | 2.5 | 2.8 | 2.4 |
| K, % | 1.0 | 1.4 | 2.0 | 2.0 | 1.2 | 2.1 | 2.41 | 1.0 |
| Mg, % | 0.44 | 0.33 | 0.40 | 0.40 | 0.29 | 0.29 | 0.28 | 0.40 |
| Na, % | 0.09 | 0.06 | 0.18 | 0.18 | 0.13 | 0.25 | 0.11 | 0.15 |
| Cl, % | 0.60 | 0.26 | 0.46 | 0.46 | 0.95 | 0.51 | 0.23 | 0.70 |
| S, % | 0.51 | 0.23 | 0.16 | 0.16 | 0.50 | 0.19 | 0.22 | 0.24 |
| Salt, % | 0.11 | 0.07 | 0.16 | 0.16 | 0.21 | 0.45 | 0.05 | 0.13 |
| DCAD, mEq/100g | -20.2 | 15.5 | 35.2 | 35.2 | -22.5 | 38.5 | 46.7 | -4.3 |
| Co, ppm | 1.3 | 1.0 | 1.0 | 1.0 | 0.68 | 0.41 | 0.34 | 0.72 |
| Cu, ppm | 17.4 | 14.9 | 13.4 | 13.4 | 4.9 | 7.3 | 2.5 | 11.7 |
| Mn, ppm | 134.7 | 104.9 | 79.0 | 79.0 | 39.7 | 23.9 | 20.0 | 80.2 |
| Zn, ppm | 137.8 | 109.5 | 75.9 | 75.9 | 53.3 | 32.1 | 26.9 | 125.4 |
| I, ppm | 1.3 | 0.9 | 0.6 | 0.6 | 0.7 | 0.4 | 0.3 | 0.8 |
| Fe, ppm | 272.4 | 261.8 | 275.3 | 275.3 | 265.2 | 190.3 | 195.6 | 276.1 |
| Se, ppm | 0.66 | 0.52 | 0.55 | 0.55 | 0.58 | 0.60 | 0.40 | 0.64 |
| Vitamin A, IU/kg | 12,809.7 | 14,124.6 | 8,293.2 | 8,293.2 | 12,684.9 | 7,935.4 | 7,535.1 | 10,732.4 |
| Vitamin D, IU/kg | 5,925.7 | 2,499.8 | 1,605.0 | 1,605.0 | 5,782.4 | 4,109.7 | 883.5 | 5,709.2 |
| Vitamin E, IU/kg | 114.0 | 120.4 | 109.4 | 109.4 | 94.4 | 82.9 | 46.5 | 103.0 |
| Moisture, % | 51.5 | 45.9 | 58.4 | 58.4 | 45.5 | 62.3 | 69.4 | 47.3 |
| Monensin, g/ton | 19.4 | 19.9 | - | - | 15.9 | 15.8 | 15.9 | 16.4 |
