## Supplementary material for "Changes in biomarkers of metabolic stress during late gestation of dairy cows associated with colostrum volume and immunoglobulin content": Table S2

**Table S2: Analyzed composition of diets.**

| | Prepartum diet | | | | | |
|---|---|---|---|---|---|---|
| | Farm A | | | | Farm B | |
| | Close up cow (n = 14) | Close up heifer (n = 13) | Far off cow (n = 12) | Far off heifer (n = 11) | Close up (n = 14) | Far off cow (n = 12) |
| DM, % | 45.9 ± 2.99 | 46.2 ± 5.64 | 43.0 ± 4.89 | 40.9 ± 4.56 | 48.8 ± 2.59 | 37.8 ± 5.8 |
| **Nutrients, DM basis (±SE)** | | | | | | |
| **Proteins** | | | | | | |
| CP, % | 12.95 ± 2.21 | 13.7 ± 0.8 | 13.74 ± 1.38 | 15.23 ± 1.67 | 13.49 ± 1.1 | 12.43 ± 2.9 |
| Adjusted Protein, % | 12.88 ± 2.22 | 13.7 ± 0.8 | 12.68 ± 1.66 | 14.29 ± 2.07 | 13.49 ± 1.1 | 11.73 ± 2.87 |
| Soluble Protein, % | 10.49 ± 15.20 | 11.85 ± 15.42 | 11.4 ± 14.29 | 14.07 ± 18.94 | 11.54 ± 16.15 | 15.91 ± 20.15 |
| Ammonia (CPE), % | 1.35 ± 0.26 | 0.89 ± 0.21 | 2.43 ± 0.35 | 2.34 ± 0.43 | 2.6 ± 0.34 | 2.68 ± 0.78 |
| ADICP, % | 1.26 ± 0.07 | 1.11 ± 0.09 | 1.98 ± 0.29 | 1.83 ± 0.24 | 1 ± 0.13 | 1.59 ± 0.33 |
| NDICP, % | 2.94 ± 0.2 | 2.55 ± 0.2 | 3.6 ± 0.7 | 3.59 ± 0.73 | 2.18 ± 0.25 | 3.07 ± 0.99 |
| **Fiber** | | | | | | |
| ADF, % | 27.1 ± 1.6 | 29.5 ± 2.89 | 39.3 ± 3.89 | 35.54 ± 2.07 | 26.72 ± 2.12 | 36.74 ± 3.77 |
| aNDF, % | 40.4 ± 3.51 | 45.58 ± 3.69 | 47.3 ± 10.39 | 46.37 ± 1.89 | 41.13 ± 3.23 | 52.81 ± 7.53 |
| Lignin, % | 4.61 ± 0.30 | 4.85 ± 0.48 | 8.74 ± 1.34 | 7.78 ± 1.19 | 4.38 ± 0.4 | 6.6 ± 1.3 |
| **Carbohydrates** | | | | | | |
| ESC-Sugar, % | 5.47 ± 1.1 | 3.81 ± 1.32 | 4.02 ± 0.35 | 3.72 ± 0.56 | 3.67 ± 0.7 | 2.36 ± 0.86 |
| Starch, % | 18.0 ± 2.27 | 19.18 ± 4.02 | 8.1 ± 2.01 | 9.87 ± 3.65 | 23.06 ± 1.87 | 11.09 ± 4.79 |
| CF, % | 2.80 ± 0.34 | 3 ± 0.6 | 3.1 ± 0.4 | 3.3 ± 0.43 | 2.66 ± 0.21 | 3.32 ± 0.42 |
| **Minerals** | | | | | | |
| Ash, % | 8.55 ± 0.53 | 7.4 ± 0.74 | 8.72 ± 1.03 | 9.29 ± 1.74 | 7.97 ± 0.67 | 9.39 ± 1 |
| Ca, % | 1.36 ± 0.31 | 0.72 ± 0.1 | 1.24 ± 0.16 | 1.27 ± 0.17 | 1.03 ± 0.27 | 0.75 ± 0.3 |
| P, % | 0.38 ± 0.04 | 0.36 ± 0.02 | 0.26 ± 0.04 | 0.3 ± 0.04 | 0.4 ± 0.04 | 0.32 ± 0.03 |
| Mg, % | 0.43 ± 0.03 | 0.36 ± 0.03 | 0.31 ± 0.06 | 0.35 ± 0.05 | 0.34 ± 0.04 | 0.27 ± 0.05 |
| K, % | 0.95 ± 0.04 | 1.19 ± 0.08 | 2.01 ± 0.16 | 1.93 ± 0.22 | 1.17 ± 0.08 | 1.98 ± 0.45 |
| S, % | 0.37 ± 0.06 | 0.23 ± 0.04 | 0.16 ± 0.03 | 0.19 ± 0.04 | 0.38 ± 0.06 | 0.19 ± 0.02 |
| Na, % | 0.08 ± 0.03 | 0.06 ± 0.02 | 0.09 ± 0.03 | 0.18 ± 0.02 | 0.11 ± 0.02 | 0.52 ± 0.17 |
| Cl, % | 0.54 ± 0.09 | 0.4 ± 0.58 | 0.47 ± 0.1 | 0.51 ± 0.17 | 0.9 ± 0.09 | 1.11 ± 0.31 |
| Fe, ppm | 398.25 ± 109.62 | 265.91 ± 44.15 | 311.5 ± 71.9 | 441.29 ± 138.36 | 371.71 ± 143.33 | 335.45 ± 123.46 |
| Mn, ppm | 126.92 ± 15.87 | 87.18 ± 31.3 | 47.3 ± 10.78 | 57.71 ± 9.72 | 75.86 ± 14.06 | 49.45 ± 7.33 |
| Zn, ppm | 118.00 ± 20.33 | 113 ± 19.61 | 36.5 ± 9.48 | 50.14 ± 11.58 | 72.21 ± 9.89 | 45.27 ± 7.32 |
| Cu, ppm | 14.92 ± 2.75 | 14.18 ± 1.72 | 11 ± 2.21 | 13 ± 4.65 | 10.71 ± 2.58 | 11.18 ± 2.79 |
| **Energy & index calculations** | | | | | | |
| TDN, % | 65.93 ± 1.93 | 65.14 ± 2.99 | 57.89 ± 1.69 | 60.26 ± 1.6 | 66.3 ± 1.87 | 59.44 ± 2.42 |
| Net Energy Lactation, Mcal/kg | 1.63 ± 0.46 | 1.61 ± 0.44 | 1.28 ± 0.04 | 1.34 ± 0.04 | 1.61 ± 0.44 | 1.32 ± 0.04 |

| | | | | | | |
|---|---|---|---|---|---|---|
| Net Energy Maintenance, Mcal/kg | 1.61 ± 0.40 | 1.57 ± 0.37 | 1.19 ± 0.04 | 1.28 ± 0.07 | 1.59 ± 0.37 | 1.23 ± 0.09 |
| Net Energy Gain, Mcal/kg | 0.95 ± 0.22 | 0.93 ± 0.18 | 0.64 ± 0.04 | 0.71 ± 0.04 | 0.95 ± 0.20 | 0.68 ± 0.09 |
| ME, Mcal/kg | 2.51 ± 0.11 | 2.49 ± 0.11 | 2.49 ± 0.88 | 2.71 ± 1.06 | 2.54 ± 0.11 | 2.47 ± 0.82 |
| Non Fiber Carbohydrates, % | 37.23 ± 4.58 | 32.54 ± 4.57 | 27.72 ± 4.88 | 28.91 ± 6.32 | 36.64 ± 3.56 | 24.57 ± 6.38 |
| Non Structural Carbohydrates, % | 24.14 ± 1.44 | 24.56 ± 2.16 | 11.92 ± 2.1 | 14.18 ± 3.35 | 26.98 ± 1.6 | 14.47 ± 3.95 |
| DCAD mEq/100g | -10.83 ± 5.63 | 12.97 ± 1.46 | 32.28 ± 4.68 | 30.94 ± 3.24 | -14.38 ± 4.89 | 28.59 ± 7.29 |

5 Results are expressed as the mean ± SE of feed samples collected every 2 weeks during sampling times. All samples were analyzed by the same external labo

6 Reflectance spectroscopy analysis and wet chemistry for minerals. Energy estimates are those predicted by the laboratory.
