## Supplementary material for "Changes in biomarkers of metabolic stress during late gestation of dairy cows associated with colostrum volume and immunoglobulin content": Table S3

**Table S3. Analytical precision biomarkers quantified using a small-scale biochemistry analyzer (CataChemWell-T, Catachem Inc., Oxford CT)**

| Analyte | Precision[1] | | | | | | | | | | | | | | | |
|---|---|---|---|---|---|---|---|---|---|---|---|---|---|---|---|---|
| | Intra-assay | | | | | | | | | | | | I | | | |
| | Sample 1 | | | Sample 2 | | | Sample 3 | | | Sample 4 | | | Sample 1 | | | |
| | Mean | SD | CV (%) | Mean | SD | CV (%) | Mean | SD | CV (%) | Mean | SD | CV (%) | Mean | SD | CV (%) | Mean |
| NEFA (m*M*) | 0.50 | 0.014 | 2.9 | 0.19 | 0.006 | 3.3 | 0.14 | 0.005 | 3.5 | 0.06 | 0.005 | 9.0 | 0.55 | 0.043 | 7.8 | 0.20 |
| BHB (mg/dL) | 16.5 | 0.24 | 1.47 | 19.8 | 0.2 | 0.99 | 9.4 | 0.15 | 1.59 | 8.7 | 0.11 | 1.26 | 16 | 0.667 | 4.18 | 19.1 |
| Cholesterol (mg/dL) | 139.4 | 1.53 | 1.1 | 47.7 | 0.73 | 1.54 | 113.5 | 5.49 | 4.84 | 123.2 | 4.34 | 3.52 | 135.2 | 8.428 | 6.23 | 46.2 |
| Glucose (mg/dL) | 42.2 | 0.89 | 2.12 | 46.5 | 1.93 | 4.16 | 52.7 | 1.6 | 3.04 | 54.9 | 1.35 | 2.46 | 41.8 | 1.994 | 4.76 | 45 |
| Total protein (g/dL) | 6.8 | 0.36 | 5.25 | 6.0 | 0.37 | 6.13 | 7.1 | 0.53 | 7.56 | 6.7 | 0.27 | 4.05 | 6.9 | 0.282 | 4.11 | 6.1 |
| BUN (mg/dL) | 14 | 0.49 | 3.45 | 13.4 | 0.36 | 2.66 | 14.0 | 0.44 | 3.13 | 15.7 | 0.33 | 2.14 | 13.8 | 0.619 | 4.49 | 13.4 |
| Albumin (g/dL) | 3.8 | 0.09 | 2.29 | 4.0 | 0.1 | 2.37 | 4.1 | 0.21 | 5.22 | 3.6 | 0.1 | 2.92 | 3.7 | 0.183 | 4.93 | 3.9 |
| Haptoglobin (g/L) | 1.55 | 0.109 | 7.10 | 0.29 | 0.019 | 6.78 | 0.72 | 0.022 | 3.09 | 0.62 | 0.025 | 4.03 | 1.54 | 0.101 | 6.54 | 0.29 |
| Calcium (mg/dL) | 8.8 | 0.08 | 0.93 | 8.2 | 0.16 | 1.89 | 8.7 | 0.23 | 2.6 | 7.9 | 0.19 | 2.43 | 8.5 | 0.411 | 4.82 | 8.2 |
| Magnesium (mg/dL) | 2.8 | 0.105 | 3.76 | 2.28 | 0.083 | 3.66 | 2.57 | 0.081 | 3.17 | 2.47 | 0.096 | 3.89 | 3.5 | 0.304 | 8.81 | 2.9 |

[1] Intra-assay precision was determined on 20 replications of four samples; inter-assay precision was determined for 15 consecutive days for three samples.
